# DISNET: A framework for extracting phenotypic disease information from public sources

**DOI:** 10.1101/428201

**Authors:** Gerardo Lagunes-García, Alejandro Rodríguez-González, Lucía Prieto-Santamaría, Eduardo P. García del Valle, Massimiliano Zanin, Ernestina Menasalvas-Ruiz

## Abstract

Within the global endeavour of improving population health, one major challenge is the increasingly high cost associated with drug development. Drug repositioning, i.e. finding new uses for existing drugs, is a promising alternative; yet, its effectiveness has hitherto been hindered by our limited knowledge about diseases and their relationships. In this paper, we present DISNET (disnet.ctb.upm.es), a web-based system designed to extract knowledge from signs and symptoms retrieved from medical databases, and to enable the creation of customisable disease networks. We here present the main features of the DISNET system. We describe how information on diseases and their phenotypic manifestations is extracted from Wikipedia, PubMed and Mayo Clinic; specifically, texts from these sources are processed through a combination of text mining and natural language processing techniques. We further present a validation of the processing performed by the system; and describe, with some simple use cases, how a user can interact with it and extract information that could be used for subsequent analyses.

## 1 Introduction

In 1796, Edward Jenner found an important link between the variola virus, which affected only humans and was highly lethal, and the bovine smallpox virus, which attacked cows and was transmitted to humans by physical contact with infected animals, and which, despite its severity, rarely resulted in death. He found that people who became infected with the latter (also called cowpox) did not subsequently catch the former; and thus, that something in the bovine smallpox virus made humans immune to variola virus. This led him to thoroughly investigate the relationship between these diseases and understand their behaviour for more than twenty years; to be finally able to find a cure for the variola virus, saving thousands of humans lives worldwide.

This discovery illustrates the importance of the knowledge that we can get from diseases and, more specifically, from how they are related. Despite the fact that in the last 200 years our understanding of diseases has greatly increased, and valuable advances have been made in this area [1], the number of those without treatment or cure is still extremely high (e.g. Alzheimer’s disease, small cell lung cancer, HIV, etc.). It is thus imperative to explore new approaches and tools to tackle them and, therefore, improve the health of the world’s population.

It is almost a truism that the search for new drugs requires a better understanding about diseases. This includes finding new insights on the relationship between diseases (which diseases are related and how), as well as the creation of public and easy-to-access large databases of diseases knowledge. During the last decade, such endeavour has been vastly facilitate by the World Wide Web. On one hand, it is possible to find free biomedical vocabularies like Unified Medical Language System (UMLS) [2], Human Phenotype Ontology (HPO) [3], [4], Disease Ontology (DO) [5] or MeSH [6], all of them offering disease classifications, disease coding standards and associated medical resources. On the other hand, one can find bioinformatic databases created by complex medical system, like DiseaseCard [7]–[9], MalaCards [10]–[12], GeneCard [13], Diseases Database (DD)1, DISEASES [14], SIGnaling Network Open Resource (SIGNOR) [15], Kyoto Encyclopedia of Genes and Genomes (KEGG) [16], MENTHA [17], PhosphositePlus [18], PhosphoELM [18], UniProtKB [19], Human Gene Mutation Database (HGMD) [20] and Comparative Toxicogenomics Database (CTD) [21]. These datasets have generally been created by processing the information from several sources, and they usually offer simple search engines; yet, not all of them have a systematic and structured form of sharing their knowledge.

It goes without doubt that the large amount of available bioinformatic resources allows both to enhance the research in the biomedical field and to have a better understanding of how the diseases behave and how can we fight them. However, most of the already mentioned sources are mainly focused on retrieving and exposing the captured knowledge and are not primarily focused on the analysis of the interactions and relationships that exists between different diseases or different disease characteristics.

In this context, several works have attempted to understand these relationships by creating and analysing disease networks. The complexity of such endeavour was soon clear, as diseases may share not only symptoms and signs, but also genes, proteins, causes and, in many cases, cures [22]–[31]. One of the most important works on the subject was published in 2007 by K.-I. Goh et al. [22], in which the HDN (Human Disease Network) was developed, a network of human diseases and disorders that links diseases based on their genetic origins and biological interactions. Different diseases were then associated according to shared genes, proteins or protein interactions. The hypothesis that different diseases, with potentially different causes, may share characteristics allows the design of common strategies regarding how to deal with the diagnosis, treatment and prognosis of a disease.

Within this line of research it is worth mentioning the Human Symptoms-Disease Network (HSDN) [23], an HDN network in which similarities between diseases were estimated through common symptoms. This is an important change in perspective with respect to previous works, in which the focus was centred on the genetic and biological origin of the diseases. In [23], diseases are defined by their clinical phenotypic manifestations, i.e. signs and symptoms; this is not surprising, as these manifestations are basic medical elements, and crucial characteristics in the diagnosis, categorization and clinical treatment of the diseases. It was then proposed to use these as a starting point to understand the existing relationships between different diseases.

Building on top of these previous works and stemming from the necessity of having exhaustive and accurate sources of disease-based information, in this paper we present the DISNET (Diseases Networks) system. DISNET aims at going one step further in improving human knowledge about diseases, not only by seeking and analysing the relations between them, but most importantly, by finding real connections between diseases and drugs, thus potentially enabling novel drug repositioning strategies.

The DISNET system allows to capture information about diseases from heterogeneous textual sources, and extract the relevant information from them as done in the HSDN work [10]. DISNET goes one step further, since among other features it is not limited to a single source of information; it provides an API-based access to the data; and integrates more powerful rules for the extraction of phenotypical manifestations from the textual sources. We also provide an evaluation of the DISNET extracted content. The captured knowledge will allow to analyse the diseases and their relationships through a network of diseases, being the current version of the system focused on phenotypical information. Future content to be introduced includes genetic and drug information to create a complex multilayer network, where each layer represents the different type of information (phenotypical, biological, drugs).

Beyond this introduction, this paper is organised as follows: Section 2 explains the technologies used in the creation of DISNET phenotypical features repository. Section 3 presents the main results obtained in the validation of the system and discussion about them, describes several simple use cases. Finally, Section 4 draws some conclusions and discusses future work.

## 2 Materials and Methods

This Section discusses the technical characteristics of the DISNET system, focusing on two aspects: the sources of information hitherto considered, and the DISNET workflow. More specifically, the last point describes how the system retrieves phenotypic information, in the form of raw texts, from the discussed sources; how these texts are processed to obtain diagnostic terms; and how these terms are validated to compile a final list of valid symptom-type terms.

### 2.1 Information Source

As it has previously been shown, it is customary for works aimed at unveiling relationships between diseases to focus on single source of information, in most cases just *abstracts* of Medline articles. On the other hand, the proposed system aims at obtaining inputs from as many sources as possible, to guarantee the recovery of as much knowledge as possible. By bringing together information from different sources, we expect them to complement each other, creating a network with a higher capacity of relating diseases. The rationale for this is that the different sources of textual knowledge, such as MayoClinic, Wikipedia, or PubMed, are written in different styles and by people with different backgrounds; the information they contain may therefore be complementary. In order to take advantage of such richness, the DISNET system allows the user to query the symptoms according to different rules: for instance, from one or multiple sources, by applying filters based on prevalence information, or on percentages of similarity among others. This clearly comes at a cost: the system should be flexible enough to be able to process sources with different structures. In the remainder of this Section we discuss the patterns used to select data sources, how they have been mined, and finally the challenges involved in such tasks.

### 2.2 Source Selection

Traditionally, in order to obtain the whole body of knowledge that mankind has accumulated about a given disease, one would refer to medical books. Although books usually contain much of the information available, they also present some important limitations: they are not constantly updated; the automatic access to their content is difficult, especially when digital versions are not available; and they are usually written for study, thus the information they contain is not structured for data mining tasks. On the other hand, one has the World Wide Web, whose main characteristic is to be (mostly) free accessible to anyone with an internet connection. It mainly offers three sources of information. Firstly, the abstract, and in some cases, the full text, of medical papers, which can be accessed through platforms like PubMed. Secondly, specialized sources of information, such as MedlinePlus, MayoClinic, or CDC. Finally, good medical data can be obtained in sources of knowledge that are not specialized, such as Wikipedia or Freebase. Note that all of them have different characteristics, in terms of comprehensiveness, degree of structure of the information, and up-to-datedness.

The criteria used for the selection of the sources of information in DISNET are: i) open access, ii) recognised quality and reliability, and iii) availability of substantial quantities of data (structured or not). This suggested to include the following three web sites in the system, which are described below: i) Wikipedia, ii) PubMed, and iii) MayoClinic. It is important to note that the system is not closed; on the contrary, thanks to its flexibility, new sources could (and will) be incorporated in the future.

### 2.3 Wikipedia

Wikipedia is an online, open and collaborative source of information. It was created by the Wikimedia Foundation and its English edition is the largest and most active one. The monumental and primary task of editing, revising and improving the quality of all articles is not performed by a core of administrators: it is instead the collaborative result of thousands of users. Consequently, this encyclopaedia is considered the greatest collective project in the history of humanity [32].

Wikipedia contains more than 155,000 articles in the field of medicine [33] and is one of the most widely used medical sources [34] by the general community [32] and also by medical specialists [35], the latter ones having deeply been involved in its enrichment [33][36]. One of the initiatives is the Cochrane/Wikipedia, which aims at increasing reliability in articles with medical content [37]. In 2014 Wikipedia was referred to as “*the single leading source of medical information for patients and health care professionals*” by the Institute of Medical Science (IMS) [38]. This stems from the fact that an increasing number of people in the medical field are becoming aware of the importance of collaborating and generating quality content in the world’s largest online encyclopaedia.

We have focused on Wikipedia in its English edition, and specifically on those articles categorized as diseases. In order to obtain a list of such articles we resort to conventional DBpedia and DBpedia-Live (DBpedia), an open and free Web repository that stores structured information from Wikipedia and other Wikimedia projects. By containing structured information, this source allows complex questions to be asked through SPARQL queries [39]. We developed a query^2^ that is able to get all the articles of Wikipedia in English referring to human diseases and run it in the **Virtuous environment SPARQL Query Editor of DBpedia**^3^. This first approach to detecting and extracting Wikipedia’s web links can be addressed in different ways and in the **Conslusions** section we will talk about them.

Even though disease articles have a standard structure, due to the very nature of Wikipedia, articles can be edited by anyone; consequently, it is possible to find articles that do not comply with the standard form that the creators of the encyclopaedia propose [40]. The structure is organized in sections, of which we have selected those whose content is related to the phenotypic manifestations of the disease. The essential sections mined by DISNET are: “*Signs and symptoms*”, “*signs and symptoms*”, “*Symptoms and causes*”, “*Signs*”, “*Symptoms*”, “*Causes*”, “*Cause*”, “*Diagnosis*”, “*Diagnostic*”, “*Causes of injury*”, “*Diagnostic approach*”, “*Presentation*”, “*Symptoms of …*“, “*Causes of …*”, and *infobox*.

The data retrieved from these sections are: i) the texts (paragraphs, lists and tables) contained in the previously described sections; ii) the links contained in these texts; and iii) the disease codes of vocabularies external to Wikipedia, which can be found in the *infoboxes* of the article. Note there are two types of *infobox*.

Fig. 2 shows an example of the external vocabulary codes retrieved in a vertical *infobox*, usually located at the beginning of the document; Fig. 1 shows an example of a horizontal *infobox*, generally located at the foot of the document. These disease codes in different vocabulary are relevant elements when searching for diseases in the system’s database. The list of external vocabularies to DISNET can be found at ^4^.

**Fig. 1.**
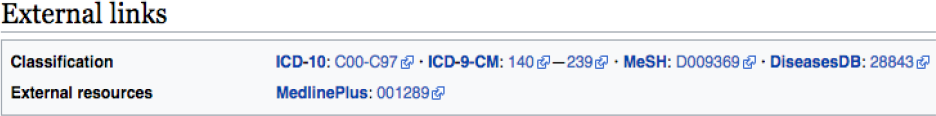
External vocabularies in a horizontal *infobox* in Wikipedia article on Cancer

**Fig. 2.**
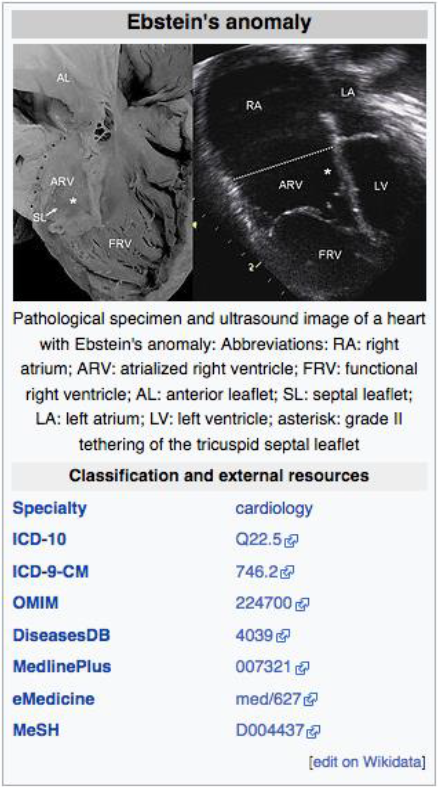
External vocabularies in a vertical infobox in Wikipedia article on Ebstein’s anomaly

### 2.4 PubMed

PubMed^5^ comprises more than 28 million biomedical literature citations from MEDLINE, life science journals and online books. Quotations may include links to full text content from PubMed Central^6^ and editorial websites [41]. As in other studies, we here only considered the abstracts of the articles, as, firstly, it is not always possible to access the full text, and secondly, the full text of articles does not follow a standard format. However, we are aware of the limitations of the extraction of information only for abstracts [42], and future versions of DISNET platform will focus in extracting the content from the full paper when possible. Note that in PubMed the information about one single disease is spread among multiple documents – as opposed to Wikipedia, in which there is a bijective relationship between articles and diseases.

Obtaining the list of diseases in PubMed involves two main steps. Firstly, one should extract the list of MeSH terms (DMTL) relating to human diseases *C*, which are categorized from *C01* to *C20* (excluding those categories such as “Animal Diseases” or “Wounds and Injuries”) as shown in the classification tree in Fig. 3^7^; and map each disease with Human Disease Ontology^8^ to obtain disease codes of the vocabulary ICD-10, OMIM, MeSH, SNOMED_CT and UMLS. Note that the use of multiple vocabularies aims at obtaining the greatest amount of means (identified codes) to identify diseases in different sources of information. As a second step, it is necessary to extract all relevant PubMed articles whose terms are associated with each of the elements of the previously extracted disease list DMTL, through PubMed’s API Entrez^9^ that we have configured to obtain, if they exist, the 100 most relevant articles of each MeSH term consulted. Specifically, for each article we retrieve: 1) abstract, 2) authors’ names, 3) unique identifier in PubMed and PubMed Central, 4) doi (digital object identifier), 5) title, 6) associated MeSH terms and 7) keywords. The workflow for extracting texts from PubMed documents is shown in Fig. 4.

**Fig. 3.**
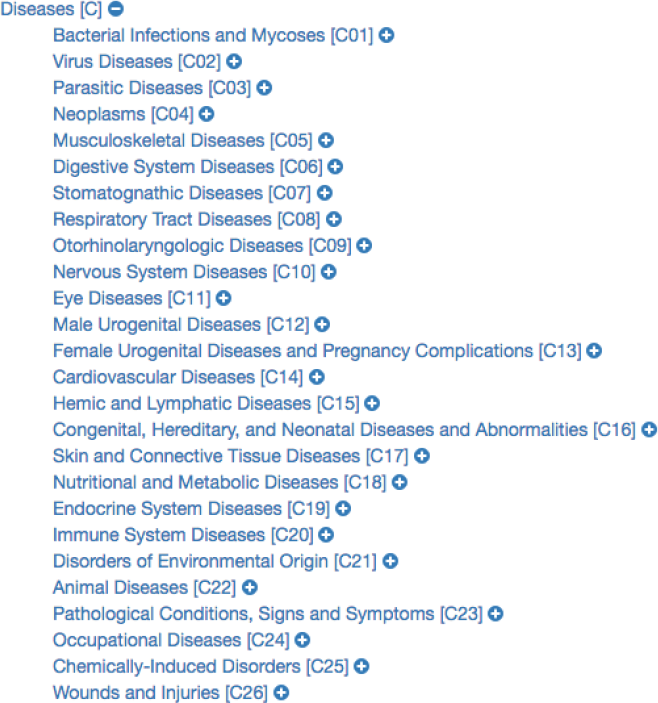
Disease MeSH Term tree clasification

**Fig. 4.**
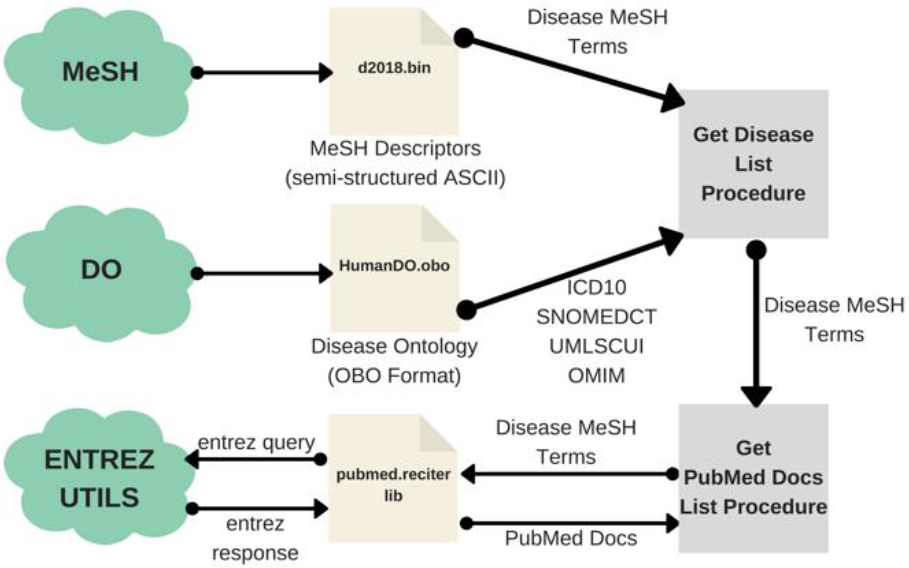
PubMed Text Extraction Procedure workflow

### 2.5 MayoClinic

According to the official website, MayoClinic^10^ is a nonprofit organization committed to clinical practice, education and research, providing expert, whole-person care to everyone who needs it. In the USA, it is considered one of the best Hospitals and Health Systems; and beyond being a provider of health services^11, 12^, it also dedicates efforts to research and publication^13^ of scientific knowledge through doctors and researchers. It thus does not come as a surprise that this online source contains relevant medical information on diseases and their phenotypic manifestations. The MayoClinic website actually contains a freely accessible list of diseases; each one of them is described in terms of an overview, the symptoms it presents, causes, diagnoses, treatments, the types of doctors who treat it and their departments or centers, among other information regarding related services provided by MayoClinic.

By early 2019 this important source of medical knowledge had 1,180 articles on diseases^14^. These articles are structured by means of sections: “Symptoms”, “Causes” and “Diagnostic”, in which we have detected a greater concentration of phenotypic textual content (paragraphs and lists).

In contrast to Wikipedia, MayoClinic does not have disease codes in external medical databases, and its list of diseases is considerably shorter; yet, it presents the advantage of being an official and curated website, being thus easier to obtain information.

### 2.6 Challenges

Mining information from the sources previously described entails several computational challenges, which may be boiled down to one requirement for the DISNET system: the need of a high versatility in data acquisition. We here review such challenges, as these partly explain the adopted software solution.

First of all, the mapping disease-webpage may take different forms. Specifically, it is one to one for Wikipedia and MayoClinic, as all the information of a disease is included in a single page; but it becomes one to many for PubMed, in which multiple articles are available for each single concept. Consulting the latter thus requires a more complex procedure.

Secondly, in most of the cases the information we want to access is always available: a user can for instance access Wikipedia or MayoClinic at any time. There are nevertheless exceptions: Freebase (which aims to be part of DISNET project in a near future) is no longer available online, and a dump has instead to be downloaded and installed locally. The system should thus be able to access both online and offline documents.

Thirdly, and as one may expect, the specific structure of each source of information is different – i.e. a page of Wikipedia has not the same structure of a PubMed article. This requires further flexibility, in terms of the development of a modular structure with specific crawlers for each source.

Finally, it is worth noting that, while here we have only considered texts, much information is available in different medias, like plain text, HTML, PDF, Word or Excel files. While not implemented at this stage, the system should be flexible enough to accommodate such sources in the future.

### 2.7 Data Retrieval and Knowledge Extraction

This section describes the general architecture of the DISNET system, including the data extraction and the subsequent knowledge extraction. In the sake of clarity, such architecture is further depicted in Fig. 5.

**Fig. 5.**
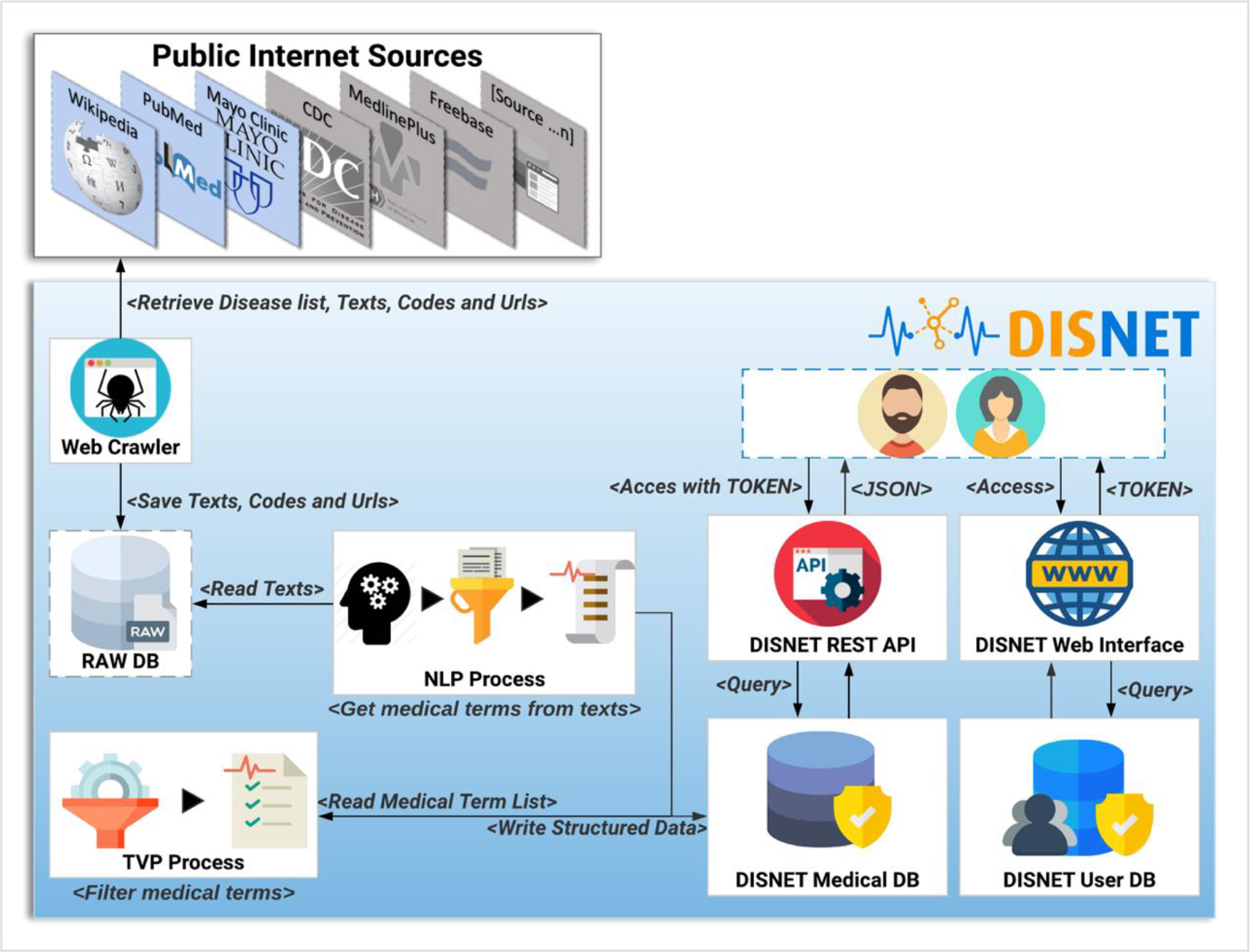
DISNET Architecture/Workflow

#### 2.7.1 The Extraction Process

The first step of the DISNET pipeline is in charge of retrieving the information from the sources previously identified and described. For each one of this, and before running the actual web crawler, the “Get Disease List Procedure” (GDLP) component is responsible for obtaining the list of diseases to be mined, thus providing links to all available disease related documents. For example, the GLDP associated to Wikipedia articles makes use of the SPARQL query^2^; similarly, the links for the PubMed’s articles are retrieved through a list of MeSH terms. However, in the case of MayoClinic, the terms are retrieved by scrapping strategies.

Once the URL list has been collected, the “Web Crawler” (WC) module is in charge of connecting to each of the hyperlinks and extracting the specific text that describes the phenotypical manifestations, as well as the links (references) contained within the texts^15^. In addition, and whenever possible, it attempts to extract information related to the coding of diseases, i.e. the codes used to identify the disease in different databases or existing data vocabularies. Currently it is able to retrieve information from more than 6,692 articles in Wikipedia, from 229,160 article abstracts in PubMed and from 1,135 articles in MayoClinic. The information mined by WC is stored in an intermediate database called “Raw DB”, which contains the raw unprocessed text.

The next step within the pipeline is called “NLP Process” (NLPP). This component is responsible for: i) reading all the texts of a snapshot, and ii) obtaining for each text a list of relevant clinical concepts/terms, discarding any unrelated paragraphs or words. At the moment NLPP uses Metamap [43][44] as a Natural Medical Language Processing tool to extract clinical terms of interest – see online NLP Tools and Configuration section^16^.

The output of the NLP process is stored in the “DISNET Medical DB” (DMDB) database. It stores, in a structured way, the medical concepts that have been obtained by the NLPP, as well as any information required to track the origin of such concepts – in order to track any error that may later be detected. Therefore, and to summarize, the information stored in a structured way in DMDB is: i) the medical concepts with their location, information and semantic types, ii) the texts from which they were extracted and the links by them contained, iii) the sections which the texts belong to, iv) the document or documents describing the disease (Web link) and v) the disease identifiers codes in different vocabulary or databases. Additional information, as the day of the extraction and the source, is further saved.

Before reaching the last step of the process, it is important to highlight the nature of the information hitherto stored. Specifically, the system has not extracted only signs or symptoms of a disease, but instead medical terms that we believe may be phenotypic manifestations of disease. It is thus necessary to filter those that are not relevant for the objective initially described.

Having clarified this, the next component of the pipeline, the “TVP Process” TVPP, reads all the concepts of a snapshot - source pair and filters them. This process is responsible for determining whether these UMLS medical terms are really phenotypic manifestations, and for storing the results back in the DMDB. TVPP is based on the Validation Terms Extraction Procedure that was developed, implemented and tested by Rodriguez-Gonzalez et al [45]. The results of this component (a purification of concepts) are thus those validated terms that we will consider as true phenotypic manifestations of diseases.

The DISNET extraction process (IEPD), i.e. the process of retrieving and storing information about diseases, basically ends here. Nevertheless, for the sake of providing an accessible and user-friendly way of retrieving and manipulating this information, DISNET also offers a REST-based interface. This is described in detail in the system website (http://disnet.ctb.upm.es/apis/disnet); also refer to Sec. 4.3 for an application example.

## 3 Results

This section describes how the medical concepts data set is built, for then validating and analysing its content. We finally present how the system could be used by means of the description of a basic use case.

### 3.1 Construction of the DB

The database in the DISNET system contains information recovered from three sources of information: Wikipedia, PubMed and MayoClinic. From Wikipedia we have 26 snapshots, from February 1^st^, 2018 to February 15^th^, 2019, for PubMed we have one snapshot, that of April 3^rd^, 2018 and for MayoClinic we have 13 snapshots, from August 15^th^, 2018 to February 15^th^, 2019. Within the system it is possible to consult, for each snapshot and source, the total number of articles with medical terms, the total number of medical terms found, the number of processed texts, the total number of retrieved codes, and the total number of semantic types found^17^.

When summing that sources, the system counts with 6,545 diseases, 2,212 medical terms from UMLS (SNOMED-CT) and 19 semantic types, which can be consulted online^18^.

Wikipedia snapshots are built using the configurations that are available online^19^. We have obtained a list of 11,074 articles catalogued as diseases in Wikipedia according to DBpedia^20^, from which we obtained 6,692 articles with at least one text referring to phenotypic knowledge of the disease, or at least one code to an external information source, 4,798 of which were found to be relevant medical concepts^21^.

The snapshot for PubMed has been built using the configuration described online^19^. This snapshot has been built on top of a list of 2,354 MeSH terms^22^ referring to human diseases, but only for 2,213 MeSH terms did we obtain information (199,013 scientific articles in total, i.e. about 0.71% of the 28 million articles existing in PubMed^23^) and of each of these PubMed articles obtained, only in 174,900 were abstracts found and only in 125,515 were relevant medical terms found. The snapshot for MayoClinic has been built on top a list of 1,188 diseases, but only on 1,125 did we obtain relevant medical terms. Fig. 6 and Fig. 7 presents some basic database statistics at an aggregated level as well as by source (for Wikipedia and PubMed). Some notable differences can be observed; for instance, the five most common terms for Wikipedia are *Pain*, *Lesion*, *Neoplasms*, *Magnetic resonance imaging*, *Inflammation* and *Malnutrition*, while for PubMed these are *Neoplasms*, *Lesion*, *Magnetic resonance imaging*, *Malnutrition* and *Inflammation*. Similarly, the three diseases with the greatest number of concepts in Wikipedia are *Kawasaki disease*, *Cerebral palsy* and *Hypoglycemia*, while for PubMed these are *Hypercalcemia*, *Cranial nerve palsy* and *Beriberi*.

**Fig. 6.**
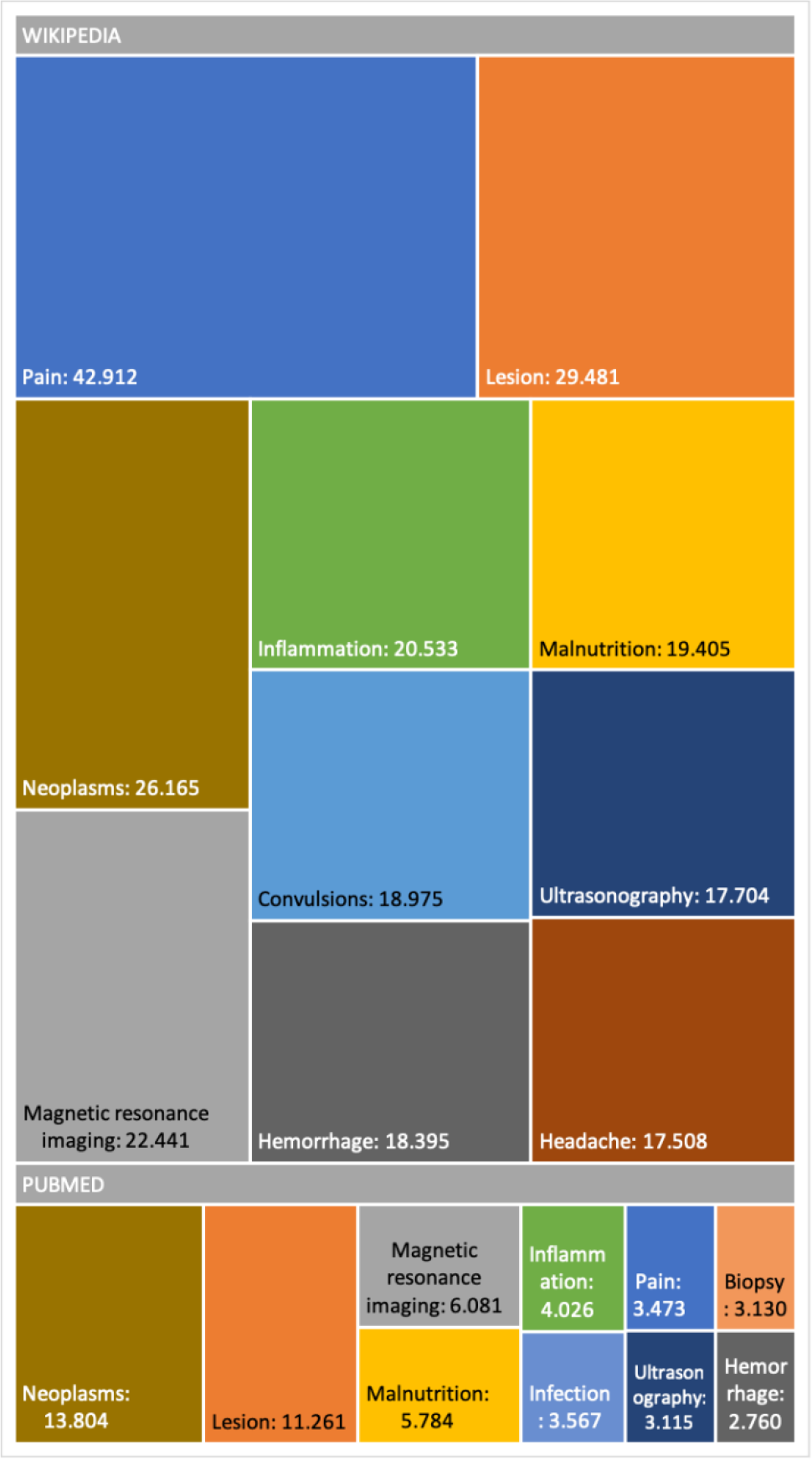
Basic database statistics (most common medical terms)

**Fig. 7.**
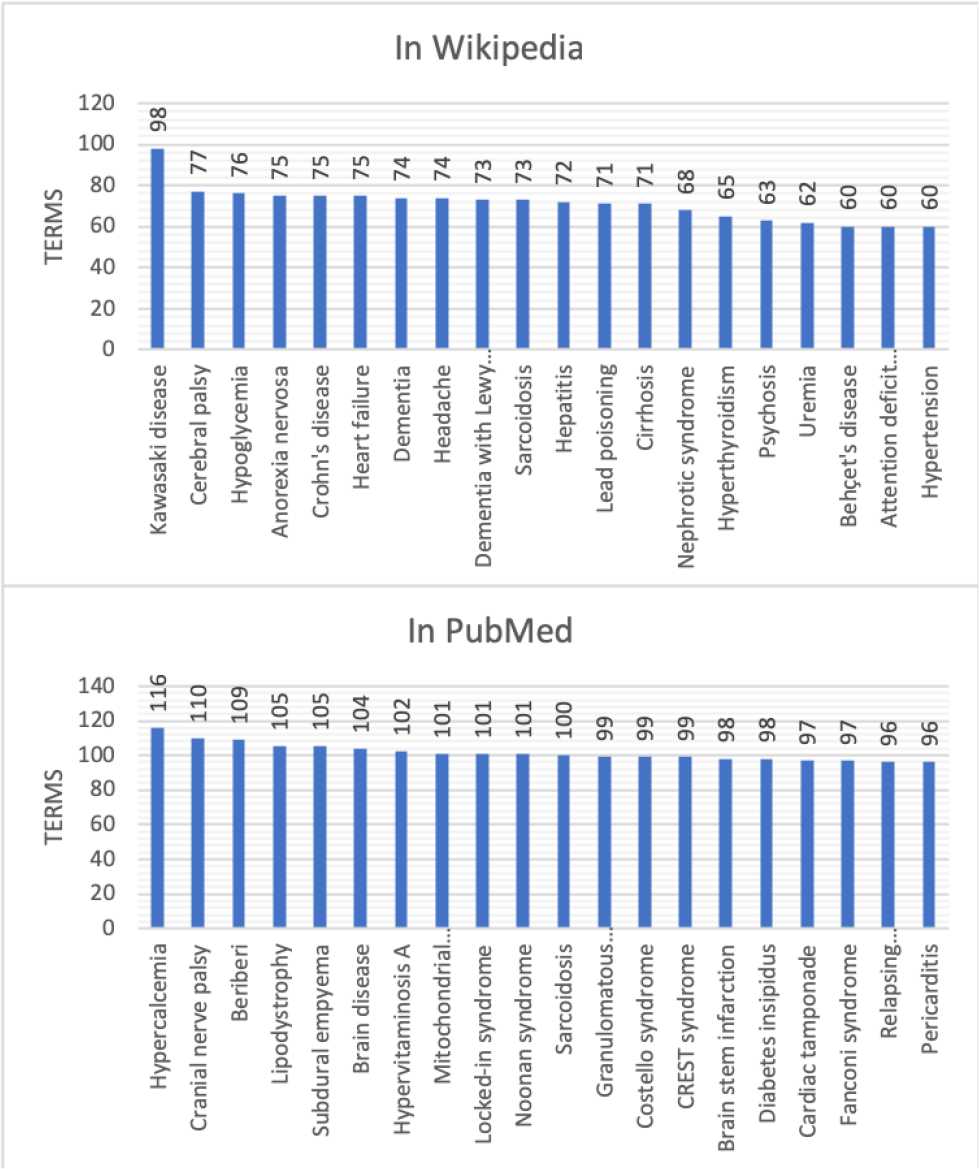
Basic database statistics (diseases with more validated medical terms. Comparison of PubMed and Wikipedia)

### 3.2 Data evaluation of the DB

In this section, we discuss the results of the validation process we executed on the system, to ensure the relevance of the diagnostic knowledge (valid medical diagnostic terms) generated through our NLP process (Metamap and TVP). The evaluation has been made on both Wikipedia and PubMed mined texts due the relevance of both sources.

The validation for Wikipedia was carried out on the February 1^st^, 2018 version, selecting 100 diseases at random with the only condition of having at least 20 valid medical terms. Similarly, the validation for PubMed has been done on the April 3^rd^, 2018 version, selecting a random sample of 100 article abstracts. These snapshots were performed at different times, and therefore with different configurations – the latter ones can be viewed online^19^. During the validation of Wikipedia, we detected that the initial configuration of Metamap did not find all the necessary medical concepts: for instance, Anxiety, Stress, Amnesia, Bulimia and other psychological concepts were missing. We therefore decided to update the initial list of semantic types to be detected (see online NLP Tools and Configuration section^16^) by adding the following elements: **Intellectual Product, Mental Process, Mental or Behavioral Dysfunction, Pathologic Function, Congenital Abnormality**.

The evaluation was conducted through a thorough manual analysis of the basic data. For each disease obtained from Wikipedia or PubMed we compared: (1) the list of medical terms extracted manually from the texts describing the disease; (2) the list of medical terms extracted by Metamap from the same texts; (3) the value (TRUE=valid or FALSE=invalid) resulting from the TVP process for each term found by Metamap; (4) the value of diagnostic relevance for a disease for each term. An example of the format of the Acute decompensated heart failure validation sheet for Wikipedia is shown in Fig. 8.

**Fig. 8.**
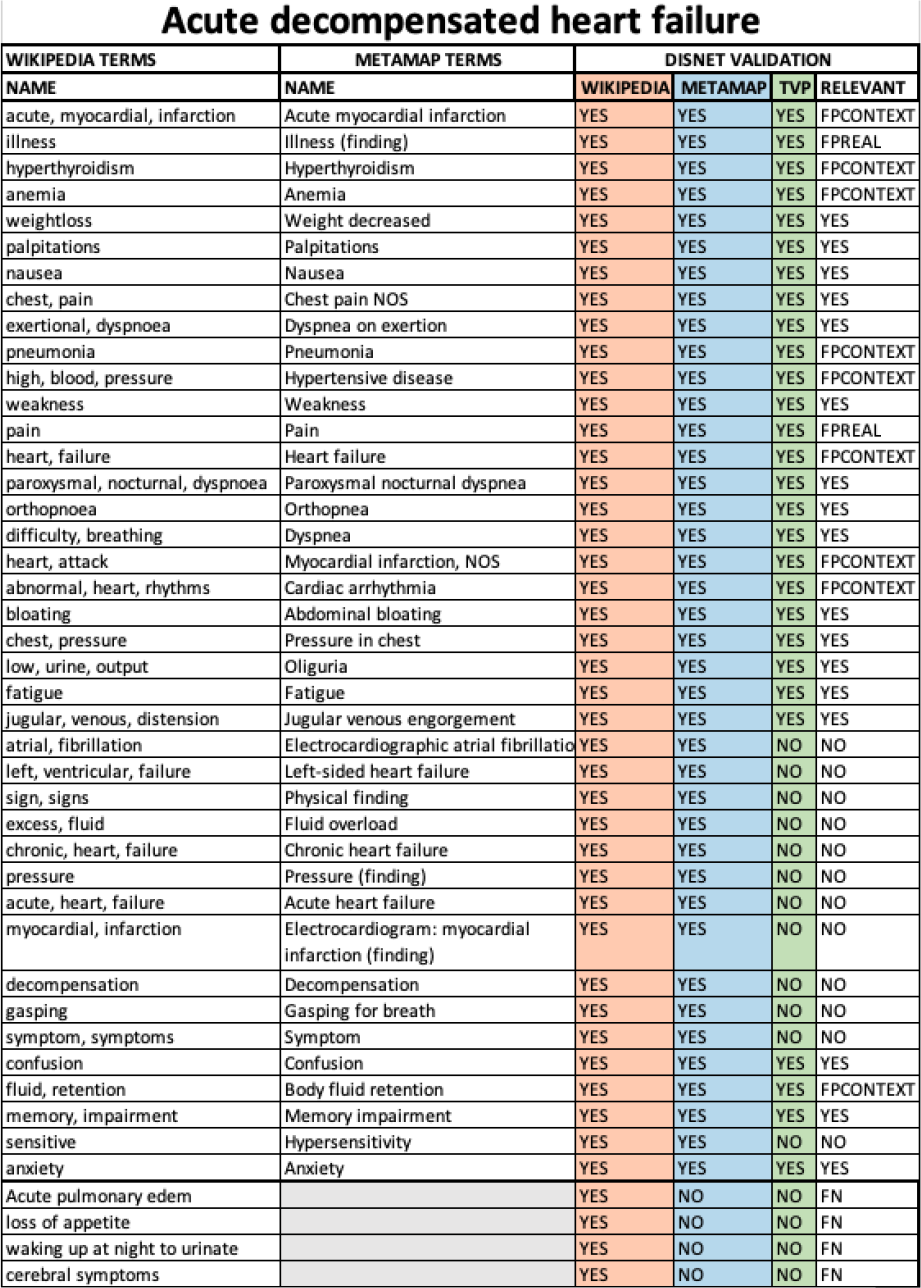
Disease Acute decompensated heart failure sheet validation from the Wikipedia snapshot of February 1^st^, 2018

It is possible to note that an additional column was also present, called RELEVANT, and which synthesises all the information available about the relevance of a term to a disease. The possible values of this column are defined as:

1. RELEVANT = **YES**. If (WIKIPEDIA = YES) & (METAMAP = YES) & (TVP = (YES or NO)), that is, it is considered to be a valid medical concept for the diagnosis of a disease.
2. RELEVANT = **NO**. If (WIKIPEDIA = YES) & (METAMAP = YES) & (TVP = NO), that is, it is considered to be a medical concept that is nonspecific, and thus too general to be helpful in the diagnosis of a disease.
3. RELEVANT = **FPREAL**. If (WIKIPEDIA = YES) & (METAMAP = YES) & (TVP = YES). The term **is not relevant** because it is considered to be a nonspecific, general concept that does not make sense for diagnosis, even though Metamap has detected it and the TVP process has evaluated it as a diagnostic term. For example, in an excerpt from Acute decompensated heart disease on Wikipedia: “*Other cardiac symptoms of heart failure include chest pain/pressure and palpitations…*”, Metamap has detected **Chest pain** and **Pain** from “*chest pain*”, both were marked as TRUE by TVP but the concept dismissed by nonspecific and general was Pain.
4. RELEVANT = **FPCONTEXT**. If (WIKIPEDIA = YES) & (METAMAP = YES) & (TVP = YES). The term **is not relevant** because it is outside the diagnostic context, even though Metamap has detected it and the TVP process has evaluated it as a diagnostic term. In other words, this term has been obtained from texts whose content is outside the diagnostic context. For example, in an excerpt from *Acute decompensated heart failure* disease on Wikipedia: “*Other well recognized precipitating factors include anemia and hyperthyroidism…*”, Metamap has detect ***Anemia*** and ***Hyperthyroidism*** which are medical terms but in context we dismiss them because they are risk factors for that disease.
5. RELEVANT = **FN**. If (WIKIPEDIA = YES) & (METAMAP = NO) & (TVP = NO). These terms were manually detected in the texts, but Metamap failed in recognising them.

The cases (3) and (4) above define situations in which the detected term is esteemed to be of no relevance, and as such represent cases of false positives. It is nevertheless necessary to discriminate the reason behind such error, which can be because: i) the term is a very general, nonspecific concept whose definition does not represent and contributes nothing to the diagnosis (FP_REAL), or ii) because the term is a medical term that is out of place with respect to the context that is narrated in the text – in other words, it could be a valid diagnostic term but not for the disease that is under validation or in the context in which have been described and therefore should be discarded (FP_CONTEXT).

Using this information for all diseases and terms, true positive (**TP**), false positive (**FP**), true negative (**TN**) and false negative (**FN**) rates were computed in order to calculate precision, recall and F1 score values as metrics to measure the performance of DISNET system. The mean values for these parameters are depicted in Fig. 9. The **TP** is all terms with (WIKIPEDIA = YES) & (METAMAP = YES) & (TVP = YES) & (RELEVANT = YES). As previously explained, the **FP** is composed of two parts, being the total FP the sum of **FP_REAL** + **FP_CONTEXT**:

**Fig. 9.**
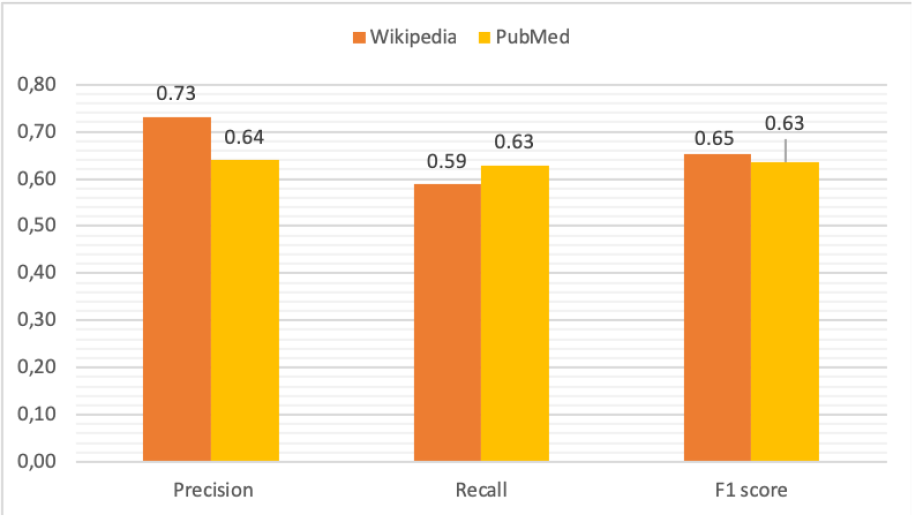
Validation metrics

- **FP_REAL** = (WIKIPEDIA = YES) & (METAMAP = YES) & (TVP = YES) & (RELEVANT = FPREAL).
- **FP_CONTEXT** = (WIKIPEDIA = YES) & (METAMAP = YES) & (TVP = YES) & (RELEVANT = FPCONTEXT).

**FN** is also composed of two parts, i.e. **FN_METAMAP** + **FN_TVP**.

- **FN_METAMAP** = (WIKIPEDIA = YES) & (METAMAP = NO) & (TVP = NO) & (RELEVANT = FN). These are terms that Metamap has not found.
- **FN_TVP** = (WIKIPEDIA = YES) & (METAMAP = YES) & (TVP = NO) & (RELEVANT = YES). These are terms that TVP has validated as false while being relevant.

Finally, the **TN** measures the TVP process (WIKIPEDIA = YES) & (METAMAP = YES) & (TVP = NO) & (RELEVANT = NO). In the Table 1 are reported the values obtained for Wikipedia and PubMed.

**Table 1.** Total values from the February 1^st^, 2018 snapshot of Wikipedia and the April 3^rd^, 2018 snapshot of PubMed

| Parameter | Wikipedia | PubMed |
| --- | --- | --- |
| TP | (31.11%) 2,075 | (31.20%) 724 |
| FP | (11.41%) 761 | (17.54%) 407 |
| FPREAL | 279 | 107 |
| FPCONTEXT | 482 | 300 |
| TN | (35.78%) 2,386 | (32.84%) 762 |
| FN | (21.68%) 1,446 | (18.40%) 427 |
| FN_METAMAP | 709 | 201 |
| FN_TVP | 737 | 226 |
| TOTAL | (100%) 6,668 | (100%) 2,320 |
| PRECISION | 0.731 | 0.640 |

Detailed results for each disease are available online, for Wikipedia^24^ and for PubMed^25^, including the list of terms manually extracted from the relevant texts of the articles, the matching with the list of terms provided by Metamap, the result of the TVP process for each term and the value of relevance as annotated by our researchers.

Results indicate that our NLP (Metamap + TVP) process is sufficiently reliable, with an accuracy of 0.731 (confidence interval of [0.710, 0.753], calculated through a Wilson’s score interval with continuity correction and a confidence level of 99%) for Wikipedia and of 0.640 (confidence interval of: [0.606, 0.680]) for PubMed (Fig. 9). The results of the calculations of these parameters for each disease can be viewed online for Wikipedia^26^ and for each abstract in PubMed^27^.

About the results for **FP** presented in Table 1, we can say that they are mainly due to the configuration used for Metamap for the extraction of terms, extended in successive extractions to avoid leaving out terms that are relevant for the detection of diseases.

Thus, one of the last extensions in the search terms added the semantic types Mental or Behavioral Dysfunction and Intellectual Product; thanks to this extension, important symptoms have been detected for certain diseases, which were not detected before, such as: *Anxiety*, *Bulimia*, *Anorexy*, *Stress*, etc. We believe that it is better to discard those terms that are not relevant than to omit those that are relevant to a disease.

It is further interesting to observe the large difference in the false positive rates between Wikipedia (11.41%) and PubMed (17.54%). We speculate that this is due to the concretion of articles. Accordingly, in Wikipedia, articles referring to one disease refer almost exclusively to that particular disease, and thus include no irrelevant terms – with a few exceptions related to differential diagnoses. Nevertheless, this is not the case of PubMed articles as a significant part of them are not so specific. Many are the articles describing real medical cases, where the symptoms are those displayed by a given patient, plus others referring to congenital diseases of the patient, or even diseases that he/she previously possessed. Consequently, the same PubMed article includes symptoms of many different diseases that, although being true medical terms and thus being recognized by Metamap, are not relevant to the disease under analysis.

For **TN**, we must also take into account that most of the terms extracted by Metamap as relevant have been purged by TVP, which has been in charge of determining which terms are relevant and which are not, so that the vast majority of terms extracted by Metamap that are not relevant to the disease have been classified in this way by TVP (35.78% for Wikipedia and 32.84% for PubMed).

In addition, we have observed that most of the true negative terms in both Wikipedia and PubMed are constant, and include: *indicated*, *syndrome*, *disease*, *illness*, *infected*, *sing*, *symptoms*, *used to*, etc.

Finally, **FN** are those terms that are relevant to the disease in question, but that have not been detected by Metamap; note that these have been manually extracted for the validation process. The vast majority of **FN** are formed by complex expressions of the language, so their detection is challenging for any NLP tool. We can further observe that the difference in the ratio of false negative between Wikipedia (21.68%) and PubMed (18.40%) is 3.28%. We believe that this difference is mainly due to the forms of expression used in both sources, with Wikipedia being more discursive, as opposed to the scientific style of PubMed.

In synthesis, we can conclude that a clear relationship can be observed between the performance of the system and the nature of the underlying data source. Specifically, while PubMed is an exclusively medical source, created, written and edited by specialists in the field, Wikipedia is a source of public information, written by anyone who has access to the web, so that the articles in it contained can be written by medical students or just users with some knowledge in the field, whose expressions cannot be assimilated to those of specialists who write PubMed. Considering that the tool used by DISNET for the extraction of medical terms (Metamap) is a medical tool, it is not surprising that it displays a greater capacity for the recognition of medical terms, as opposed to more colloquial terms formed by more complex phrases; thus, there are terms such as “*Swollen lymph glads under the jaw*”, or “*sensation of swelling in the area of the larynx*”, that Metamap cannot recognize.

### 3.3 A use case

To illustrate the possible use of the DISNET system, we here present a simple use case, which consists of the creation of several basic DISNET queries, and the visualization of the corresponding results.

#### 3.3.1 Creation of DISNET queries

For the sake of simplicity, we will here focus on two of the most important characteristics of DISNET: **i)** the ability to create relationships between diseases according to their phenotypic similarity (**C1**) and **ii)** the ability to increase/improve the phenotypic information of diseases by means of periodic extractions of knowledge (**C2**).

The scenario C1 implies obtaining data for two diseases, which we suspect may share symptoms; we will here use “Influenza” and “Gastroenteritis”. The resulting DISNET queries are:

1. disnet.ctb.upm.es/api/disnet/query/disnetConceptList?source=wikipedia&version=2018-08-15&diseaseName=Influenza&matchExactName=true
2. disnet.ctb.upm.es/api/disnet/query/disnetConceptList?source=pubmed&version=2018-04-03&diseaseName=Influenza&matchExactName=true
3. disnet.ctb.upm.es/api/disnet/query/disnetConceptList?source=wikipedia&version=2018-08-15&diseaseName=Gastroenteritis&matchExactName=true
4. disnet.ctb.upm.es/api/disnet/query/disnetConceptList?source=pubmed&version=2018-04-03&diseaseName=Gastroenteritis&matchExactName=true

We have here used the DISNET query “**disnetConcepList**”, which allows retrieving the list of “**DISNET Concepts**” associated with a given disease. The parameters of this query include: “**diseaseName**”, with the name of the disease; “**matchExactName**”, to indicate that the search by disease name is exact; and “**source**” and “**snapshot**”, to respectively indicate the source and snapshot we want to consult. In this case, we selected to consult the two sources Wikipedia and PubMed, and respectively the snapshots of August 15^th^, 2018 and April 3^rd^, 2018. Note that the result will consists of four total lists, two for each disease. To illustrate, Fig. 11 shows an extract of the response from the query (3).

As for the scenario C2, it requires retrieving data for a disease whose list of symptoms may have changed with time, i.e. either increased or decreased. As an example, we considered the disease “Acrodynia”, and executed the following DISNET queries:

1. disnet.ctb.upm.es/api/disnet/query/disnetConceptList?source=wikipedia&version=2018-02-01&diseaseName=Acrodynia&matchExactName=true
2. disnet.ctb.upm.es/api/disnet/query/disnetConceptList?source=wikipedia&version=2018-02-15&diseaseName=Acrodynia&matchExactName=true

Note that, as in C1, we have here used the query “**disnetConceptList**”; nevertheless, we have here executed it twice, on the same disease (**Acrodynia**) and two different snapshots: February 1^st^, 2018 and February 15^th^, 2018.

#### 3.3.2 Visualization of the result of the DISNET queries

Once the results of the query have been retrieved, the next natural step is their visualization; while the actual output format may vary according to the needs of each specific project, for the sake of clarity we here created a graph representation by using the external tool Cytoscape^28^. In both scenarios (i.e. C1 and C2) we generated relationships between diseases and their symptoms, with the aim of visualizing the value and scope of the medical data stored and processed by DISNET. In Fig. 10.b we see the relationship between the Influenza and Gastroenteritis diseases on one hand (highlighted in white rectangles), and the set of symptoms on the other. Symptoms were obtained from two different sources, specifically Wikipedia and PubMed: relationships are then respectively represented by red and blue edges. Common symptoms are merged by the layout algorithm in the center of the graph; the medical terms that are not common among the two diseases, on the contrary, form a peripheral shell. Note that “**Influenza**” has 59 DISNET Concepts and “**Gastroenteritis**” has 47, 19 of which are in common.

**Fig. 10.**
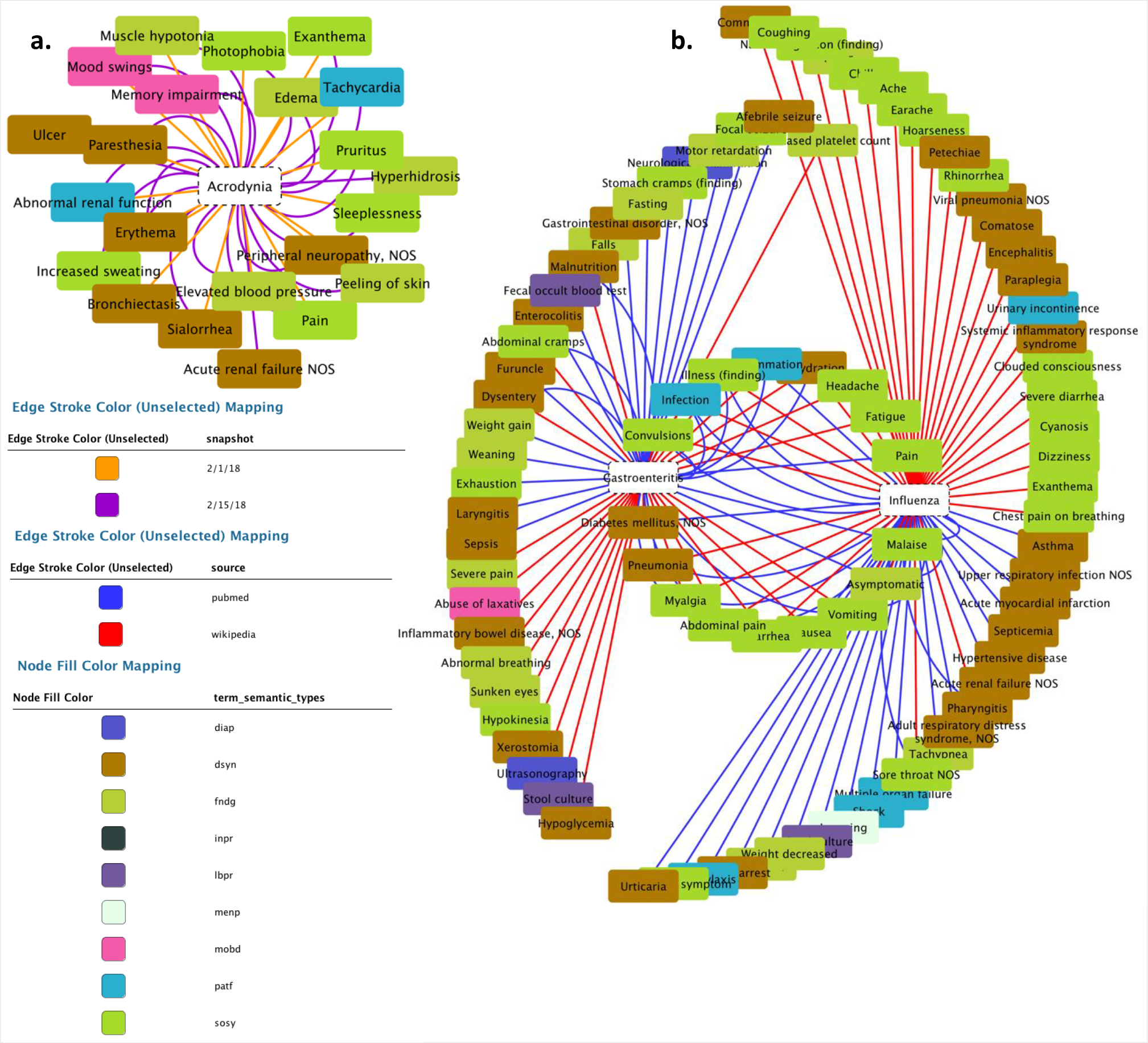
a) Network of graphs representing the evolution of phenotypic knowledge in Wikipedia and b) Network of graphs representing similar medical terms between two diseases.

**Fig. 11.**
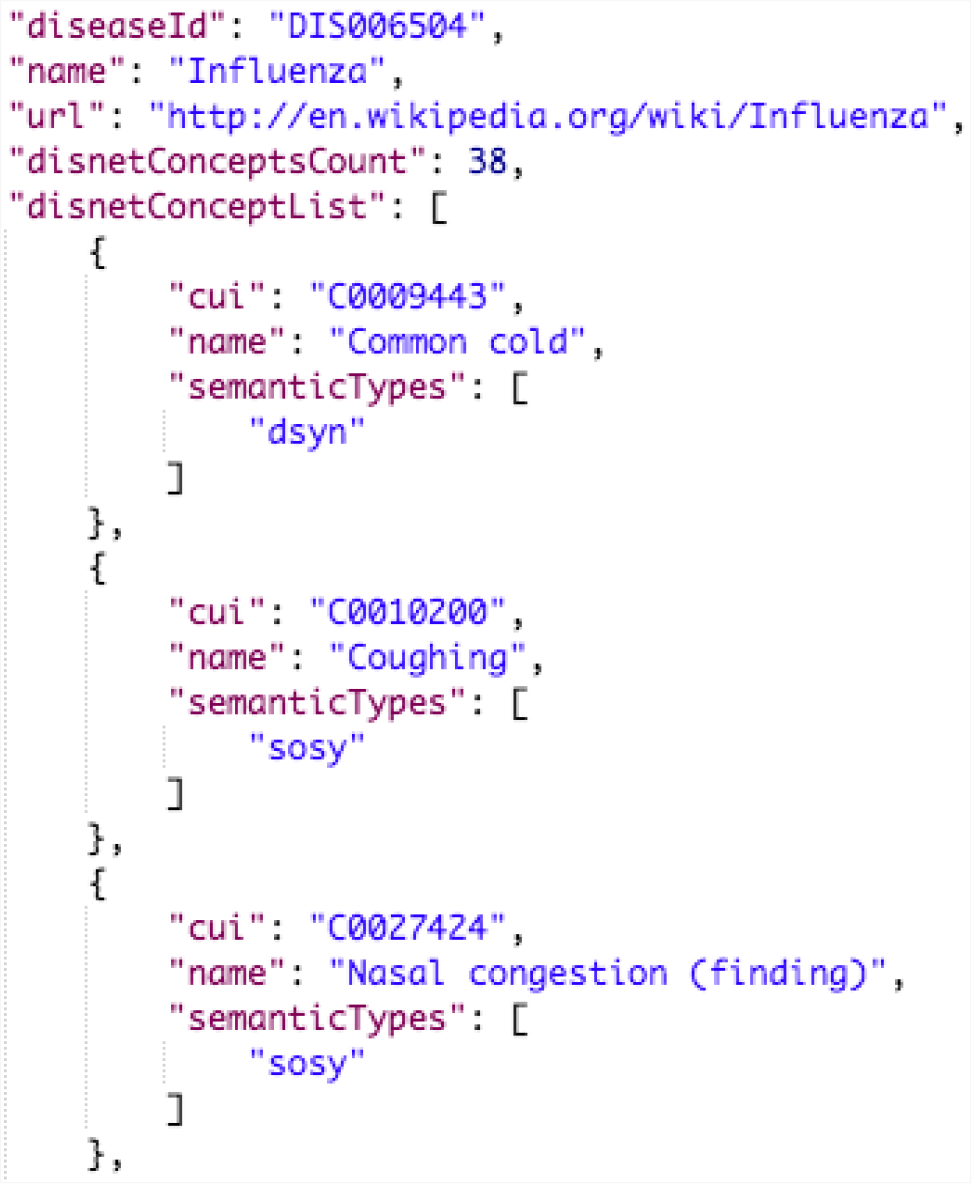
Answer to the DISNET query “disnetConcepList” C1.(1)

In Fig. 10.a we observe the network representation of the disease “**Acrodynia**” and of its 18 medical terms, 15 of which were found in the snapshot of February 1^st^, 2018 and three new ones in that of February 15^th^, 2018. This is thus an example of an increase in phenotypic knowledge.

This simple use case illustrates how the DISNET system allows generating a network of diseases and their symptoms on a large scale, and that it provides the right environment to know how diseases are related according to their phenotypic manifestations. By applying similarity algorithms, such as Cosine [23][46][47] or the Jaccard index [48], it is possible to estimate the similarity between two diseases, and thus to focus further medical analyses on those pairs showing a large overlap. These features will be also implemented as native features in next DISNET release.

## 4 Conclusions

This work presented the DISNET system, starting from its underlying conception, up to its technical structure and data workflow. DISNET allows retrieving knowledge about the signs, symptoms and diagnostic tests associated with a disease. It is not limited to a specific category (all the categories that the selected sources of information offer us) and clinical diagnosis terms. It further allows to track the evolution of those terms through time, being thus an opportunity to analyse and observe the progress of human knowledge on diseases. We also presented the DISNET REST API, which aims at sharing the retrieved information with the wide scientific community. We further discussed the validation of the system, suggesting that it is good enough to be used to extract diseases and diagnostically-relevant terms. At the same time, the evaluation also revealed that improvements could be introduced to enhance the system’s reliability.

Among the potential lines of future works, priority will be given to increasing the number of information sources, by including other websites like Medline Plus or CDC. Secondly, we are considering the possibility of extending the TVP procedure, by adding new data sources, with the aim of increasing the number of validation terms and hence of reducing the number of false negatives. Note that this could also partly be achieved by resorting to a different NLP tool to process the input texts, as for example to Apache cTakes [49]. Future implementations of DISNET also aim to provide ways to automatically compute the similarity between diseases (by using already mentioned and well-known similarity metrics), extending the DISNET platform to include biological and drug information and developing new visualization strategies, among others.

1 http://www.diseasesdatabase.com

2 https://midas.ctb.upm.es/gitlab/disnet/paperdisnet/blob/master/get_diseases_query.sparql

3 https://dbpedia.org/sparql

4 https://midas.ctb.upm.es/gitlab/disnet/paperdisnet/blob/master/wikipedia_medical_vocabularies.txt

5 https://www.ncbi.nlm.nih.gov/pubmed/

6 https://www.ncbi.nlm.nih.gov/pmc/

7 https://b.nlm.nih.gov/treeView

8 http://www.obofoundry.org/ontology/doid.html

9 https://www.ncbi.nlm.nih.gov/home/develop/api/

10 https://www.mayoclinic.org

11 https://www.mayoclinic.org/es-es/about-mayo-clinic/quality/rankings

12 https://www.mayoclinic.org/es-es/about-mayo-clinic/office-diversity-inclusion

13 https://www.mayo.edu/research/publications

14 https://www.mayoclinic.org/es-es/diseases-conditions

15 https://jsoup.org/

16 http://disnet.ctb.upm.es/apis/disnet#NLP_Tools_and_Configuration

17 https://midas.ctb.upm.es/gitlab/disnet/paperdisnet/blob/master/knowledge_sources

18 https://midas.ctb.upm.es/gitlab/disnet/paperdisnet/blob/master/DISNET_summing_source_counts

19 https://midas.ctb.upm.es/gitlab/disnet/paperdisnet/blob/master/snapshot_settings.txt

20 https://midas.ctb.upm.es/gitlab/disnet/paperdisnet/blob/master/wikipedia_diseases_articles_by_dbpedia.txt

21 https://midas.ctb.upm.es/gitlab/disnet/paperdisnet/blob/master/wikipedia_articles_with_relevant_terms.txt

22 https://midas.ctb.upm.es/gitlab/disnet/paperdisnet/blob/master/mesh_terms_human_diseases.txt

23 https://midas.ctb.upm.es/gitlab/disnet/paperdisnet/blob/master/list_pubmed_papers.txt

24 https://midas.ctb.upm.es/gitlab/disnet/paperdisnet/tree/master/wikipedia_validation_sheets

25 https://midas.ctb.upm.es/gitlab/disnet/paperdisnet/tree/master/pubmed_validation_sheets

26 https://midas.ctb.upm.es/gitlab/disnet/paperdisnet/blob/master/wikipedia_individual_validation_results.csv

27 https://midas.ctb.upm.es/gitlab/disnet/paperdisnet/blob/master/pubmed_individual_validation_results.csv

28 http://www.cytoscape.org

